# Development of Systemic Immune Dysregulation in a Rat Trauma Model with Biomaterial-Associated Infection

**DOI:** 10.1101/2020.01.10.901769

**Authors:** Casey E. Vantucci, Hyunhee Ahn, Mara L. Schenker, Pallab Pradhan, Levi B. Wood, Robert E. Guldberg, Krishnendu Roy, Nick J. Willett

## Abstract

Orthopedic biomaterial-associated infections remain a large clinical challenge, particularly with open fractures and segmental bone loss. Invasion and colonization of bacteria within immune-privileged canalicular networks of the bone can lead to local, indolent infections that can persist for years without symptoms before eventual catastrophic hardware failure. Host immunity is essential for bacterial clearance and an appropriate healing response, and recent evidence has suggested an association between orthopedic trauma and systemic immune dysregulation and immunosuppression. However, the impact of a local infection on this systemic immune response and subsequent effects on the local response is poorly understood and has not been a major focus for addressing orthopedic injuries and infections. Therefore, this study utilized a model of orthopedic biomaterial-associated infection to investigate the effects of infection on the long-term immune response. Here, despite persistence of a local, indolent infection lacking outward symptoms, there was still evidence of long-term immune dysregulation with systemic increases in MDSCs and decreases in T cells compared to non-infected trauma. Further, the trauma only group exhibited a regulated and coordinated systemic cytokine response, which was not present in the infected trauma group. Locally, the infection group had attenuated macrophage infiltration in the local soft tissue compared to the non-infected group. Our results demonstrate widespread impacts of a localized orthopedic infection on the systemic and local immune responses. Characterization of the immune response to orthopedic biomaterial-associated infection may identify key targets for immunotherapies that could optimize both regenerative and antibiotic interventions, ultimately improving outcomes for these patients.

## INTRODUCTION

Severe musculoskeletal trauma of the extremities is a major cause of death and disability in both military and civilian populations with over 12 million fractures annually in the United States (1). Despite advances in fracture stabilization and wound management, morbidity and complication rates remain high with up to 10% of patients experiencing delayed or compromised healing (2). The most common complications include infections and delayed or non-unions, resulting in longer rehabilitation times for patients and increased treatment costs (3). In particular, implant-associated infections occur in 0.5-5% of orthopedic implants and can lead to impaired bone healing and hardware failure, requiring subsequent interventions, revision surgeries, and long-term antibiotics, increasing total health and societal costs (4,5).

In open fractures, orthopedic hardware and biomaterial implants provide necessary fracture stability and support tissue regeneration; however, they also provide an ideal environment for bacteria to colonize and grow. *Staphylococcus aureus* is the most frequent orthopedic-implant associated organism, found in 34% of orthopedic infections (6). *S. aureus* can adhere to biomaterial implant surfaces within hours and form a complex structure surrounded by self-generated extracellular polymeric substance matrix, called a “biofilm,” which is comprised of proteins, polysaccharides, lipids, and nucleic acids (7,8). The secreted proteins from bacterial cells within the biofilm enhance immune evasion and promote *S. aureus* infiltration into the canalicular network within the bone (9). Invasion and colonization of the canalicular network makes the infection even more challenging to treat as the immune system tries to clear the infection that is inaccessible deep within these networks (10). Successful survival of the bacteria in the host can result in subsequent bone lysis with decreased osteoblast viability and increased bone resorption resulting from both bacterial factors and host inflammation in response to the implant being recognized as a foreign body (11,12). Patients with biomaterial-associated infections will often require additional surgeries and aggressive antimicrobial therapy which is estimated to cost as high as $150,000 per patient (13).

In some cases, biomaterial-associated infections are indolent and remain local, meaning that the infection is slow-growing and does not result in a systemic response or pose an immediate threat to the patient. In more severe cases, local biomaterial-associated infections that trigger acute innate and adaptive immune responses may lead to a systemic spread of the bacterial infection, bacteremia, subsequent pro-inflammatory systemic immune responses, and the onset of sepsis (14–16). Sepsis is marked by a hyper-inflammatory phase and cytokine storm that results in a high fever, heart rate, and breathing rate, and if left unchecked, can even lead to multiple organ failure and death. Improvements in patient management and supportive care have drastically improved patient survival through this hyper-inflammatory phase which is subsequently followed by an immunosuppressive phase. Following the cytokine storm, there is a marked decrease in host immunity including apoptosis-induced loss of innate and adaptive immune effector cell populations as well as an increase in immune suppressor cell populations, including T regulatory cells (Tregs) and myeloid-derived suppressor cells (MDSCs) (17–20). With improvements to patient care, sepsis has transitioned from a hyper-inflammatory disorder to an immunosuppressive disorder, with almost 70% of deaths occurring after the initial inflammatory phase due to the subsequent impaired immunity that make patients more vulnerable to infection recurrence or nosocomial infections (21). Current treatment strategies for sepsis are now not only aimed at treating the initial systemic infection but also at preventing cellular apoptosis and boosting host immunity (22–25).

Interestingly, a similar systemic immunosuppressive response has been observed clinically in a subset of severe trauma patients without associated infection and has been termed “sterile sepsis” due to similarities in the innate and adaptive immune responses but a lack of bacterial presence (26). Systemic immune dysregulation and immunosuppression following severe trauma has also been associated with increased susceptibility to infection and poor healing outcomes, especially for more severe injuries (27–29). Immediately after injury, massive early innate immune responses trigger a systemic inflammatory response syndrome (SIRS), resulting in over-production of pro-inflammatory mediators (e.g. Type I Interferons, IL-1, IL-6, IL-8, TNFa) and systemic activation of innate immune cells (30–32). Concurrently, the body’s defense mechanisms trigger a compensatory anti-inflammatory response syndrome (CARS) through systemic up-regulation of immunosuppressive mediators (e.g. Il-10, TGF-b) and cells, such as Tregs and MDSCs (28,33). A balance between the SIRS and CARS responses leads to restoration of systemic immune homeostasis typically within the first week or two following injury and is associated with successful healing and regeneration. However, when the CARS response overwhelms the initial SIRS response, systemic immune dysregulation and immunosuppression develops, marked by increases in immunosuppressive mediators in response to low-level persistent inflammation caused by damage-associated molecular patterns (DAMPs) at the injured site (34). Patients exhibiting systemic immune dysregulation following non-infected trauma are more susceptible to opportunistic infections, have an increased risk for complications, and exhibit impaired healing and regeneration (27–29). The immune evasion and immunosuppressive mechanisms of *S. aureus* in biomaterial-associated orthopedic infections may further exacerbate conditions in trauma patients with systemic immune dysregulation; however, the mechanisms and development of this response are poorly understood.

Host immunity is essential for bacterial clearance and appropriate and regulated healing; however, the role of the systemic immune response is not well understood and has not been a major focus for addressing orthopedic injuries and infections thus far. Orthopedic biomaterial infections are associated with immune evasion mechanisms that alter immune responses and long-term systemic immune responses to trauma have more recently been associated with immune dysregulation, immune suppression, and low-level persistent inflammation. This could create a complex and poorly understood immune environment that may contribute to inadequate healing and infection recurrence in patients who have developed biomaterial-related infections after trauma. In this study, the objective was to characterize the systemic immune response to an orthopedic biomaterial-associated infection following severe trauma. We hypothesized that a local, indolent infection combined with trauma would lead to long-term systemic immune dysregulation and immunosuppression. A better understanding of the immune response to severe trauma with associated infection may be essential for optimizing therapeutic interventions and improving outcomes for these patients.

## METHODS

### Micro-organism preparation

A bio-luminescent strain of *Staphylococcus aureus* (Xen29, PerkinElmer, Waltham, MA) was cultured in Luria Bertani (LB) medium containing 200 μg/ml kanamycin at 37°C, under aerobic conditions while agitated at 200 rpm for ∼2-3 hours.

### Surgical procedures

All animal care and experimental procedures were approved by the Veterans Affairs Institutional Animal Care and Use Committee (IACUC) and carried out according to the guidelines. Unilateral 2.5mm femoral segmental defects were created in 21-week old female Sprague-Dawley rats (Charles River Labs) in a similar manner to previous segmental defects (35). Briefly, an anterolateral incision was made along the length of the femur and the vastus lateralis was split with blunt dissection. A modular fixation plate was affixed to the femur using miniature screws (JI Morris Co., Southbridge, MA, USA). The 2.5mm segmental defect was then created in the diaphysis using a Gigli wire saw (RISystem, Davos, Switzerland). A collagen sponge with (collagen + infection, n=6) or without (collagen only, n=7) bacteria inoculum (*S. aureus* at 10^7^ CFU) was placed in the defect (Figure 1A). The fascia was then sutured closed with absorbable 4-0 sutures, and the skin was closed with wound clips. Buprenorphine SR (0.03 mg/kg; 1 ml/kg) was used as an analgesic and applied via subcutaneous injection. Body temperature and weight were recorded prior to surgery and monitored longitudinally after surgery at days 1, 3, 7, 14, 28, and 56.

**Figure 1.**
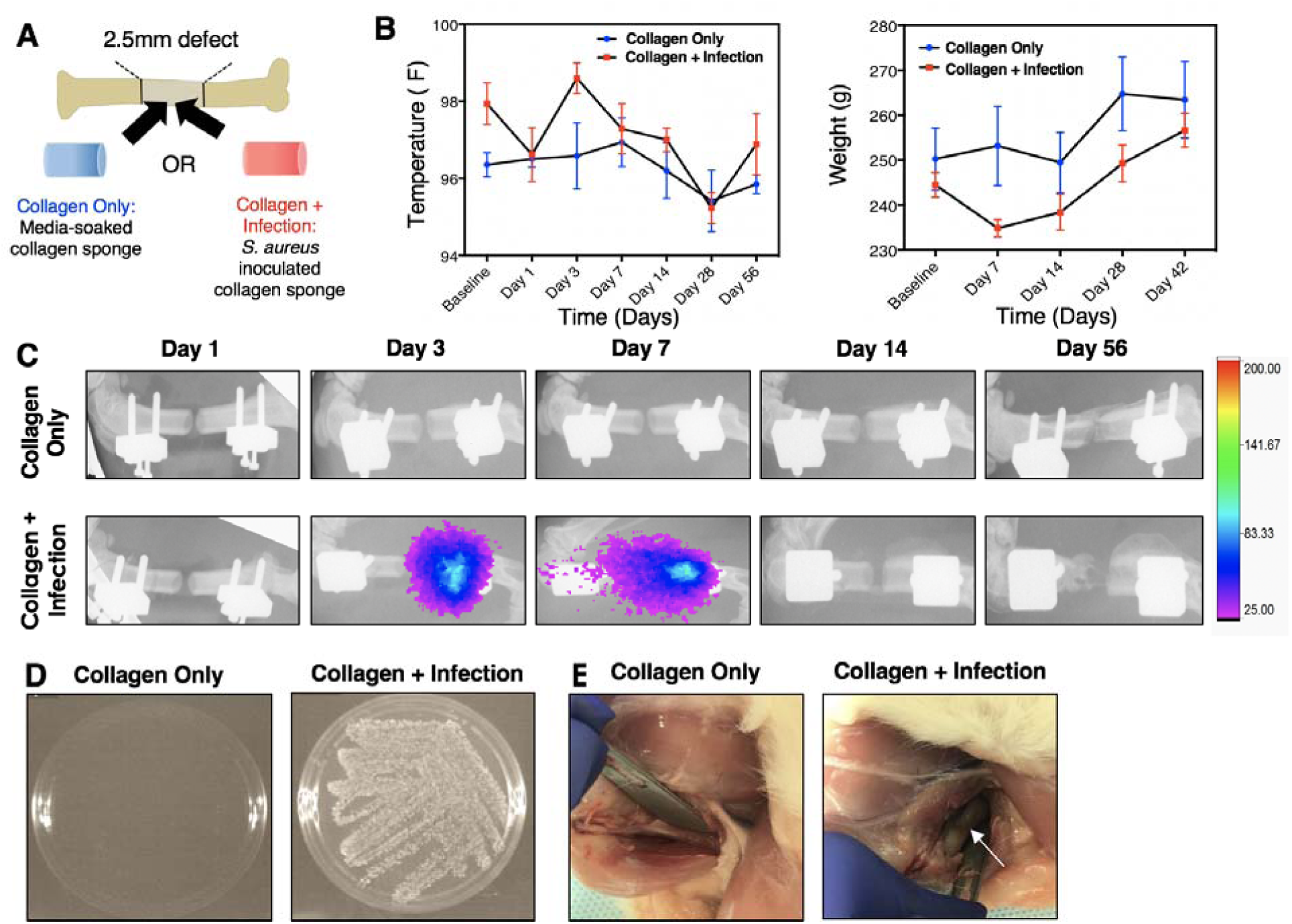
A) Each animal will receive a 2.5mm segmental bone defect supported by an internal fixation plate. An untreated, media-soaked collagen sponge or a collagen sponge inoculated with *S. aureus* will be placed into the defect site prior to closure of the surgical site. B) Temperature (left) and weight (right) of the collagen only, non-infected animals and the collagen + infection animals. No significant differences between groups overall or at any time point for temperature and weight were observed according to repeated measures 2-way ANOVA. C) Serial bioluminescent and radiograph images taken at Days 1, 3, 7, 14, and 56. Bioluminescent signal appeared in infection animals at Day 3 post-surgery and was present up to Day 7. D) Bacterial culture following wound swab of non-infected and infected animals at the Day 56 endpoint. Bacterial growth is present in the collagen + infection group, but not in the collagen only group. E) Representative images of the thighs of euthanized non-infected and infected animals. The white arrow points to gray necrotic soft tissue and purulence around the hardware.

### Microbiological analysis

Bacterial metabolic activity was monitored *in vivo* using serial bioluminescent (BL) scanning (In-Vivo Xtreme, Bruker Corp., Billerica, MA, USA) at Days, 3, 7, 14, and 56. X-rays were taken together with BL scanning as reference images. Bacterial contamination was also confirmed at 8 weeks post-surgery via wound swab culture.

### Immune Characterization

#### Circulating cellular analysis

Whole blood was collected via the rat tail vein longitudinally at days 0 (baseline), 1, 3, 7, 14, 28, and 56 for flow cytometry analysis. Red blood cells were lysed using 1X RBC lysis buffer (eBioscience) according to the manufacturer’s instructions. Cells were then fixed using Cytofix fixation buffer (BD) and resuspended in buffer containing 2% fetal bovine serum (FBS) in 1X PBS and stored at 4°C until stained. Prior to staining, cells were blocked with purified anti-rat CD32 (BD) to prevent nonspecific binding. Cells were then stained for various immune cell populations, including T cells (CD3+) and T cell subsets (CD4+, CD8+, and FoxP3+), B cells (B220+), and MDSCs (His48+CD11b+) with specific anti-rat antibodies (eBioscience). Sample data were collected using a BD Accuri C6 flow cytometer and analyzed with FlowJo. Gates were positioned with less than 1% noise allowed based on fluorescent minus one (FMO) controls.

#### Tissue cellular analysis

At the week 8 endpoint, tissues were harvested for immune cell population analyses including: local soft tissue adjacent to the defect site, the spleen, and bone marrow from both the contralateral leg and the tibia from the injured leg. Cells were stained for various immune cell populations, including B cells (B220+), MDSCs (His48+CD11b+), tissue macrophages (His36+), and hematopoietic stem cells (HSCs, CD45+CD90-) in the bone marrow only with specific anti-rat antibodies (eBioscience). Staining and analysis procedures were the same as for the circulating cellular analyses.

#### Cytokine and chemokine analysis

Serum was isolated from whole blood at the same timepoints by allowing it to clot overnight at 4°C. Samples were then centrifuged at 1500g for 10 min and the supernatant was collected and stored at −80°C until analysis. Multiplexed chemokine and cytokine analysis was performed using Milliplex MAP Rat Cytokine/Chemokine Magnetic kit (Millipore Sigma) and analyzed using a MAGPIX Luminex instrument (Luminex). Median fluorescent intensity values with the background subtracted were used for multivariate analyses.

### Micro-computed tomography

Bone mineralization from the injured site was quantitatively assessed using micro-computed tomography (uCT) scans (Micro-CT40, Scanco Medical, Bruttisellen, Switzerland) at 8 weeks. Samples were scanned with a 20 µm voxel size at a voltage of 55 kVp and a current of 145 µA. The bone volume was quantified only from the defect region, and new bone formation was evaluated by application of a global threshold corresponding to 50% of the cortical bone density (386 mg hydroxyapatite/cm^3^).

### Histological analysis

After euthanasia at 8 weeks post-surgery, upper hindlimb explants were harvested and fixed with 10% neutral buffered formalin (10% NBF) for 3 days and then stored in 70% ethanol until processing. To observe mineralized bone structure, decalcified bone samples were embedded in paraffin and cut using a microtome (Leica Microsystems, Wetzlar, Germany) to an average thickness of 10 µm. Deparaffinized slides were then stained with Hematoxylin &Eosin (E&O) staining to demonstrate new bone formation. Images were obtained with an Axio Observer Z1 microscope (Carl Zeiss, Oberkochen, Germany) and captured using the AxioVision software (Carl Zeiss, Oberkochen, Germany).

### Statistical analysis

Statistical significance for quantitative results were assessed using appropriate parametric or non-parametric tests. For data that met the assumptions, an unpaired Student’s t-test or analysis of variance (ANOVA) with repeated measures was used. Multiple comparisons were made using Sidak’s multiple comparisons test, and significance was determined by p values less than 0.05. For data that did not meet the assumptions for parametric tests, the Mann-Whitney test was used. Additionally, mixed effects modeling (REML) was used for repeated measures analysis of the circulating immune cell data. Numeric values are presented as the mean ± SEM. All statistical analysis was performed using GraphPad Prism 8.0 software (GraphPad, La Jolla, CA, USA).

Luminex data was further analyzed by partial least square discriminate analysis (PLSDA) in MATLAB (Mathworks) with the partial least squares algorithm by Cleiton Nunes (available on the Mathworks File Exchange) following z-scoring to normalize the data. PLSDA analysis reduces the dimensionality of the input variables into a set of latent variables (LVs) that maximally separate discrete groups (i.e. collagen + infection versus collagen only). Latent variables are composed of profiles of the input variables that represent their relative contributions to the latent variables, and thus the separation between the groups. Monte Carlo sub-sampling with 1,000 iterations was done to determine the standard deviation for each of the individual signals in the LV loading plot. For each iteration, all but one of the samples were randomly sampled and a new PLSDA model was determined. Sign reversals were corrected by multiplying each sub-sampled LV by the sign of the scalar product of the new LV and the corresponding LV from the total model. The mean and standard deviation were then computed for each signal across all iterations. Lastly, heat maps of the z-scored data for all cytokine values were generated.

## RESULTS

### Establishment of a local infection associated with a biomaterial implant

#### Temperature and weight

Body temperature and weight change were measured longitudinally after surgery. Neither body temperature nor weight were significantly different between the infection animals and the non-infected, collagen only animals at any time point or overall (Figure 1B). Throughout the 56-day time period, infection animals showed normal weight gain and did not have elevated body temperatures compared to the collagen only animals, indicating that the infection did not spread systemically.

#### Bioluminescent scans and endpoint evaluation of infection

Bioluminescent scans indicating metabolic activity of the bacteria were assessed following inoculation (Figure 1C). Bioluminescent signal was present in infection animals as early as Day 3 post-surgery and up until Day 7; however, it should be noted that bioluminescence is indicative of metabolic activity of the cells, not necessarily cell presence. No signal was observed in collagen only animals at any time point. Despite the absence of bioluminescent signal beyond Day 7 *in vivo*, wound swab culture conducted at the Day 56 endpoint confirmed the presence of bacteria in the infection animals (Figure 1D). No bacterial growth was observed in culture following wound swab of the non-infected, collagen only animals. Further, inspection of the thighs of euthanized infection animals showed gray, necrotic soft tissue and purulence around the implant hardware, which was not present in collagen only animals (Figure 1E).

### Bone regeneration and histological analysis

At the Day 56 endpoint, bone explants were harvested for micro-computed tomography (uCT) and histological evaluation. There were no significant differences in new bone formation between the collagen only and infection animals, although the collagen only group did have higher peak bone formation (Figure 2A). Further, histological analysis revealed an abnormal ectopic periosteal response in the bone from the infection animals, with aberrant and scattered periosteal bone formation (Figure 2B). Non-infected bone showed new bone formation at the defect region according to hematoxylin and eosin staining.

**Figure 2.**
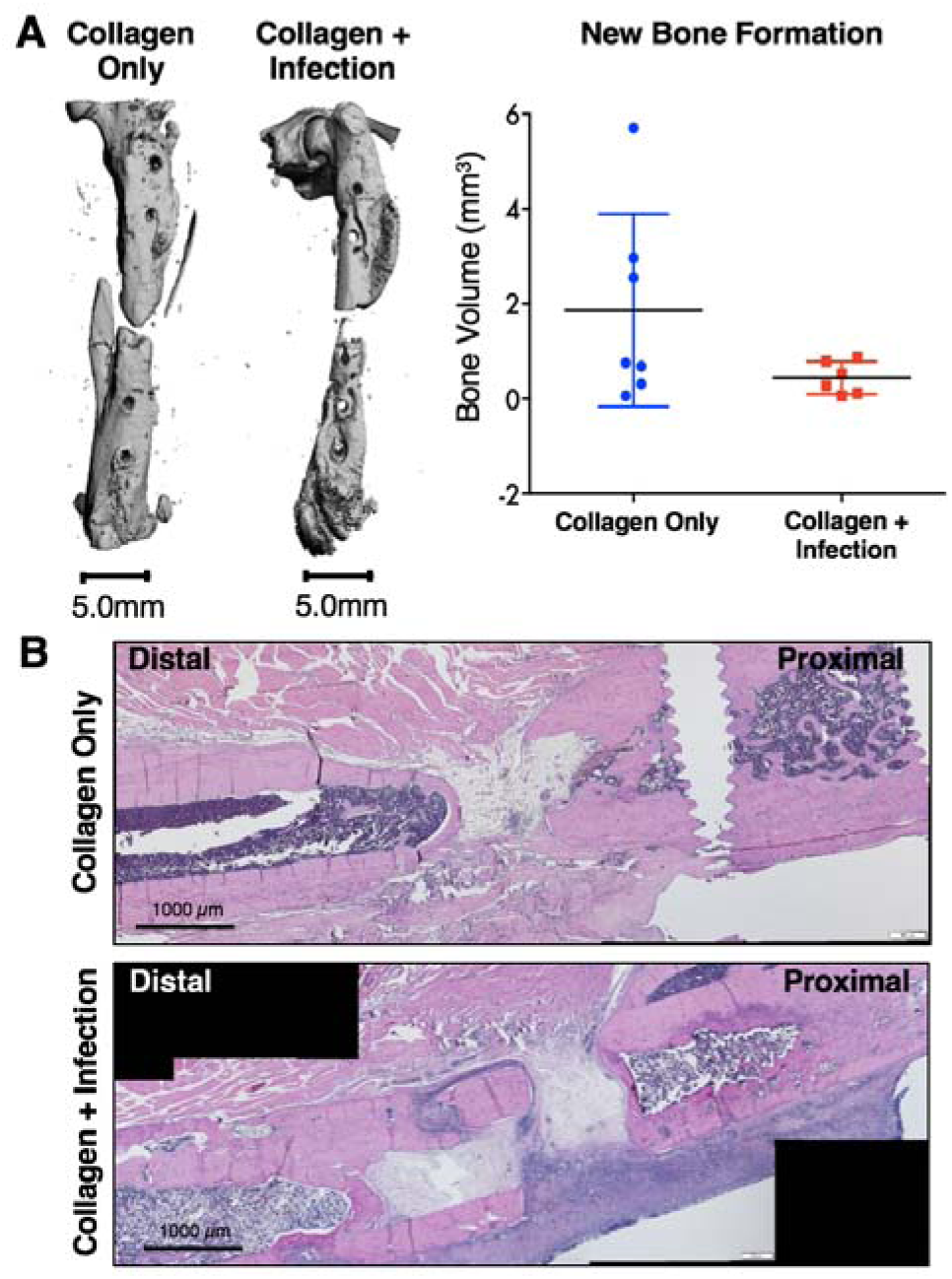
A) Representative endpoint (Day 56) uCT reconstructions (left). Quantification of uCT scans showed no significant differences between the collagen only and collagen + infection group via Mann-Whitney test (p=0.2343). B) Representative Hematoxylin &Eosin (E&O) stains of the defect depicting bone formation (pink) and cell presence (nuclei stained purple) within the defect region.

### Local infection alters local and systemic immune profiles

#### Systemic immune cell populations

Circulating immune cell populations of both immune effector cells and immunosuppressive cells were evaluated using flow cytometry and revealed differences in the systemic immune response in the infection versus collagen only animals (Figure 3). Immune effector cells evaluated included both T and B cells (Figure 3A), as well as the helper T cell and cytotoxic T cell subsets (Figure 3B). Immunosuppressive cells evaluated included MDSCs and Tregs (Figure 3C), an immunoregulatory T cell subset. At Day 1 post-injury, there was a significant decrease in circulating T cells, helper T cells, and cytotoxic T cells in both the infection and collagen only groups compared to the baseline. Over time, T cell populations, including helper T cell and cytotoxic T cell subsets, gradually increased in both groups until Day 28. However, while the T cell populations in the collagen only group increased back to baseline levels, the T cell populations in the infection group remained significantly lower overall compared to baseline and the collagen only group. For B cells, there was a similar decrease in cell numbers in the collagen only group, whereas there was an increase in B cells in the collagen + infection group, which peaked at Day 7. Following Day 7, B cells in the infection group continue to decline, whereas B cells in the collagen only group remained relatively constant. For the immunosuppressive cell types, at Day 1 post-injury, there was a significant systemic increase in MDSCs in both the infection and collagen only groups compared to baseline with the highest peak level of MDSCs in the infection group. In contrast, MDSCs in both groups gradually decreased until Day 28; however, MDSCs in the infection group overall remained elevated (p = 0.06) compared to the collagen only group. Tregs showed no significant differences between the groups. Overall, there were decreases in T cells, helper T cells (p=0.05), and cytotoxic T cells (p=0.06) and increases in immunosuppressive MDSCs (p=0.06) in the infection group compared to the collagen only group.

**Figure 3.**
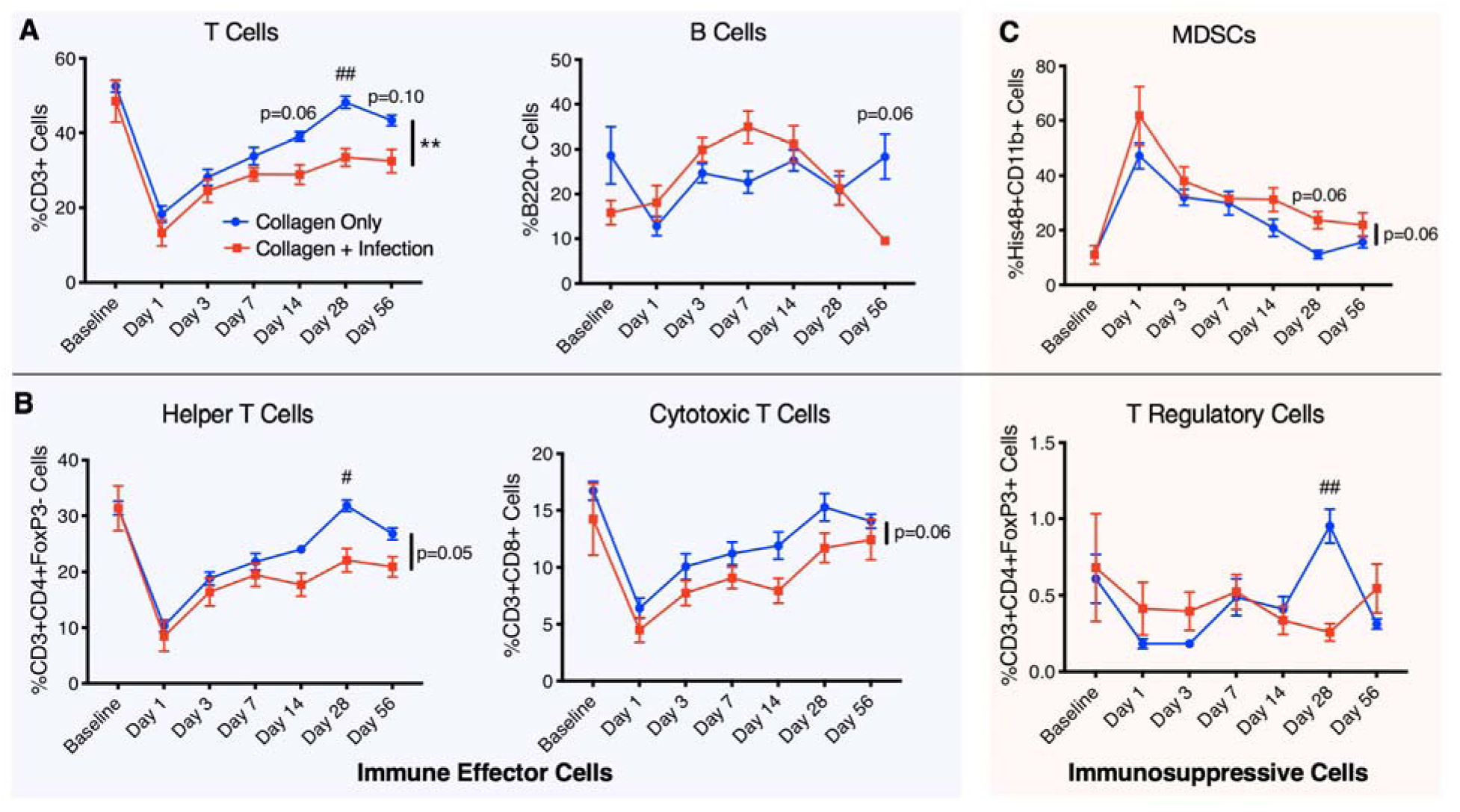
Longitudinal analysis of immune cells circulating in the blood including A) lymphocytes (T cells and B cells), B) immunosuppressive MDSCs, and C) T cell subsets that include helper T cells, cytotoxic T cells, and T regulatory cells. These cell types are divided into immune effector cells (T cells, B cells, helper T cells, and cytotoxic T cells) and immunosuppressive cells (MDSCs and T regulatory cells). Overall differences between groups are indicated with a line and the p value or by a ** (p<0.01). Differences between groups at specific timepoints are indicated with the p value or by a # (p<0.05) or ## (p<0.01). P values were determined by fitting a mixed-effects model (REML) with Sidak’s multiple comparisons test, an analysis similar to repeated measures 2-way ANOVA that can handle missing data points.

#### Tissue immune profiles

At the Day 56 endpoint, tissue was collected from the spleen, the soft tissue adjacent to the defect, and the bone marrow in the injured and contralateral legs. Similar to systemic cellular analysis, immunosuppressive MDSCs were elevated in infection animals compared to collagen only animals in both local soft tissue and the bone marrow, but not in the spleen (Figure 4). Macrophages, another immune effector cell type, were found to be decreased in the infected group in the local soft tissue (adjacent to the bone defect), the spleen, and the bone marrow compared to the collagen only group. Additionally, B cells in the local soft tissue and the spleen were found to be lower in the collagen + infection group compared to the collagen only group. Analysis of hematopoietic stem cells in the bone marrow revealed a slight decrease in the infection animals compared to the collagen only animals but were not significantly different (Figure 4C,D). Despite a decrease in circulating T cells in infection animals compared to collagen only animals, there were no significant differences in T cell populations in any of the tissues.

**Figure 4.**
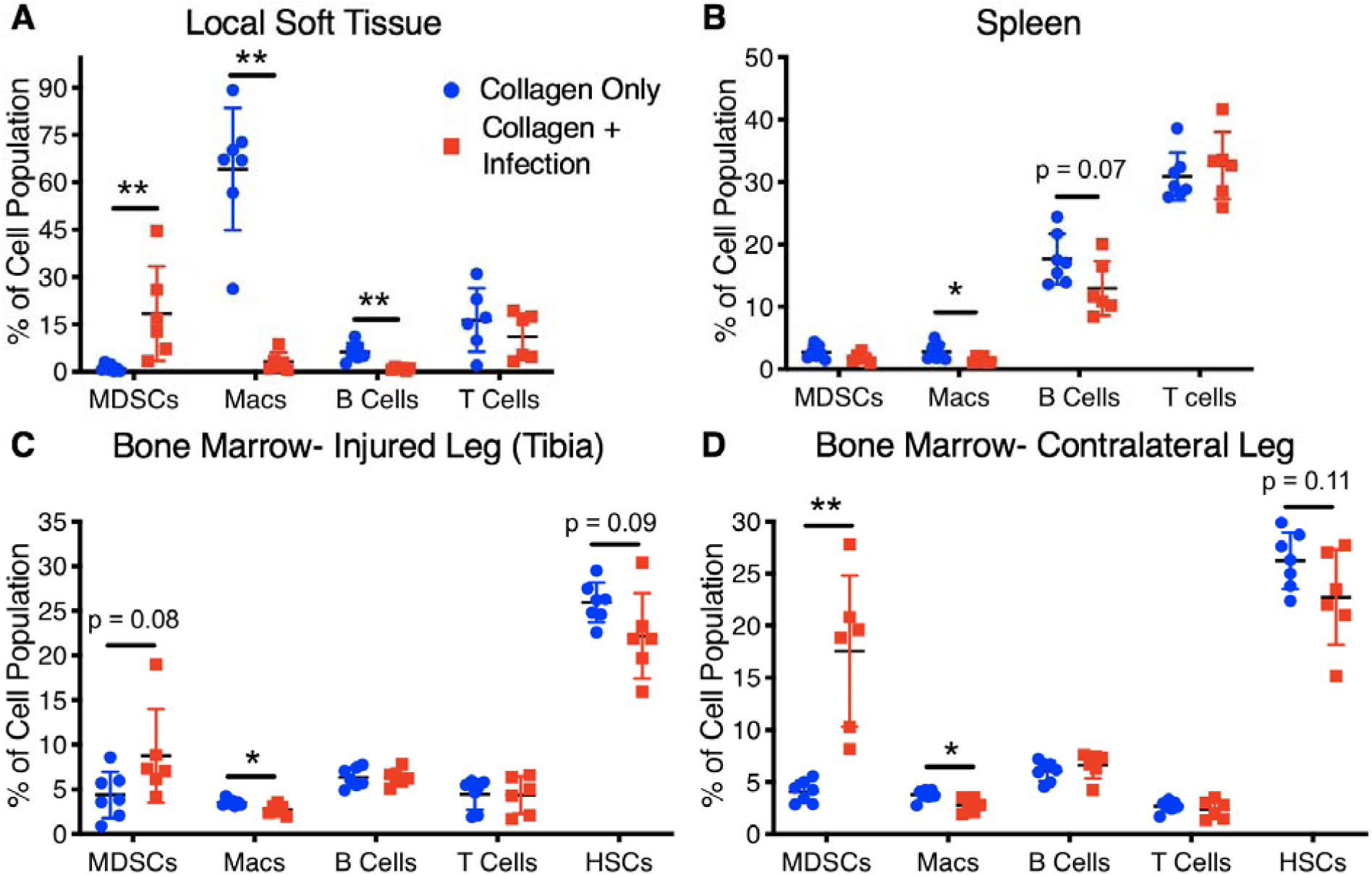
Endpoint cellular analyses (Day 56) of tissues including the local soft tissue adjacent to the defect, the spleen, the bone marrow from the contralateral leg, and the bone marrow from the tibia of the injured leg. Differences between groups are indicated by a p value or by * (p<0.05) or ** (p<0.01). P values obtained using Student’s t test or non-parametric Mann-Whitney test when variances are significantly different between groups.

#### Cytokine and chemokine analysis

Cytokine and chemokine data were z-scored (mean subtracted and normalized to standard deviation for each cytokine) and plotted on a heat map over time (Figure 5A). The collagen only group revealed coordinated up-regulation of specific cytokines at Day 1, Day 7, and Day 14, which appeared to resolve by Day 28. Qualitatively, the coordinated regulation of cytokine and chemokine levels was not observed in the infection group. Quantitatively, there were no significant differences in baseline levels between the groups for any of the cytokines (Supplementary Figure 1A). At Day 1, endothelial growth factor (EGF), macrophage chemoattractant protein-1 (MCP-1), and LIX were significantly upregulated in the collagen only group compared to the infection group (Figure 5B). All of these factors are involved in cell migration and recruitment, cell survival, and differentiation. Additionally, IL-2, a cytokine important for T cell function, was upregulated in the infection group compared to the collagen only group at Day 1 (Figure 5B). No other cytokines were significantly different between groups at Day 1 (Supplementary Figure 1B). On Day 3, RANTES, a chemokine important for leukocyte recruitment, was significantly upregulated in the collagen only group compared to the infection group; however, IL-10, a general immunosuppressive cytokine, was significantly elevated in the infection group compared to the collagen only group (Figure 5C). No other cytokines were significantly different between groups at Day 3 (Supplementary Figure 1C). By Day 7 and 14, a large number of both pro- and anti-inflammatory cytokines were all significantly upregulated in the collagen only group compared to the infection group (Supplementary Figure 1D,E). The inflammation in the collagen only group resolved between Day 14 and Day 28, with cytokine and chemokine levels returning to baseline levels. At Day 28, there were no significant differences between groups for any of the cytokines (Supplementary Figure 1F); however, by Day 56, several cytokines including IL-4, IL-12p70, IL-17A, and TNFa were all upregulated in the infection group compared to the collagen only group (Supplementary Figure 1G).

**Figure 5.**
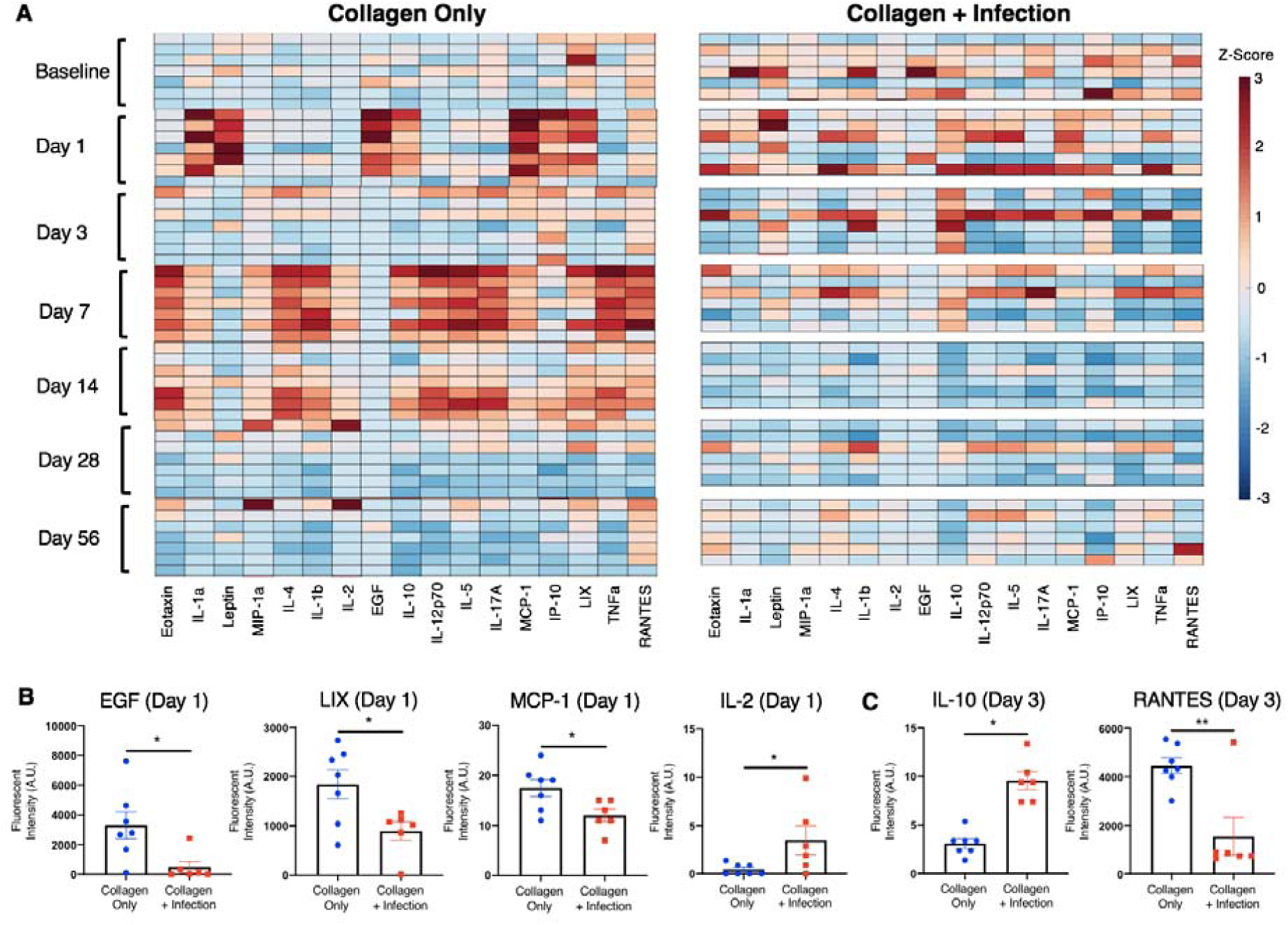
A) Heat map of z-scored cytokine levels (each column represents a different cytokine) in the collagen only group (n=7, one n per row) and the collagen + infection group (n=6, one n per row) prior to surgery (Baseline) and at Days 1, 3, 7, 14, 28, and 56 post-surgery. Cytokine levels at the two earliest time points that exhibited significant differences between groups are shown for B) Day 1 and C) Day 3. Significance was determined using Student’s t-test or with Mann-Whitney test for non-parametric data with p<0.05 (*) and p<0.01 (**).

#### Overall systemic immune characterization

To investigate the overall contributors that distinguished the infected group from the non-infected group, partial least squares discriminant analysis (PLSDA) was conducted on data aggregated from all timepoints post-injury, including cellular and chemokine data (Figure 6A). This analysis revealed a significant separation of the two groups according to latent variable 1 (LV1) (Figure 6B). The LV1 loading plot shows the major contributors to positive LV1 values (collagen only group) and negative LV1 values (collagen + infection group) (Figure 6C). Here, the main contributors to the collagen only group were T cells and the T cell effector subsets, including helper T cells and cytotoxic T cells. Contributing most to the infection group were the immunosuppressive MDSCs, B cells, and the immunosuppressive cytokine IL-10.

**Figure 6.**
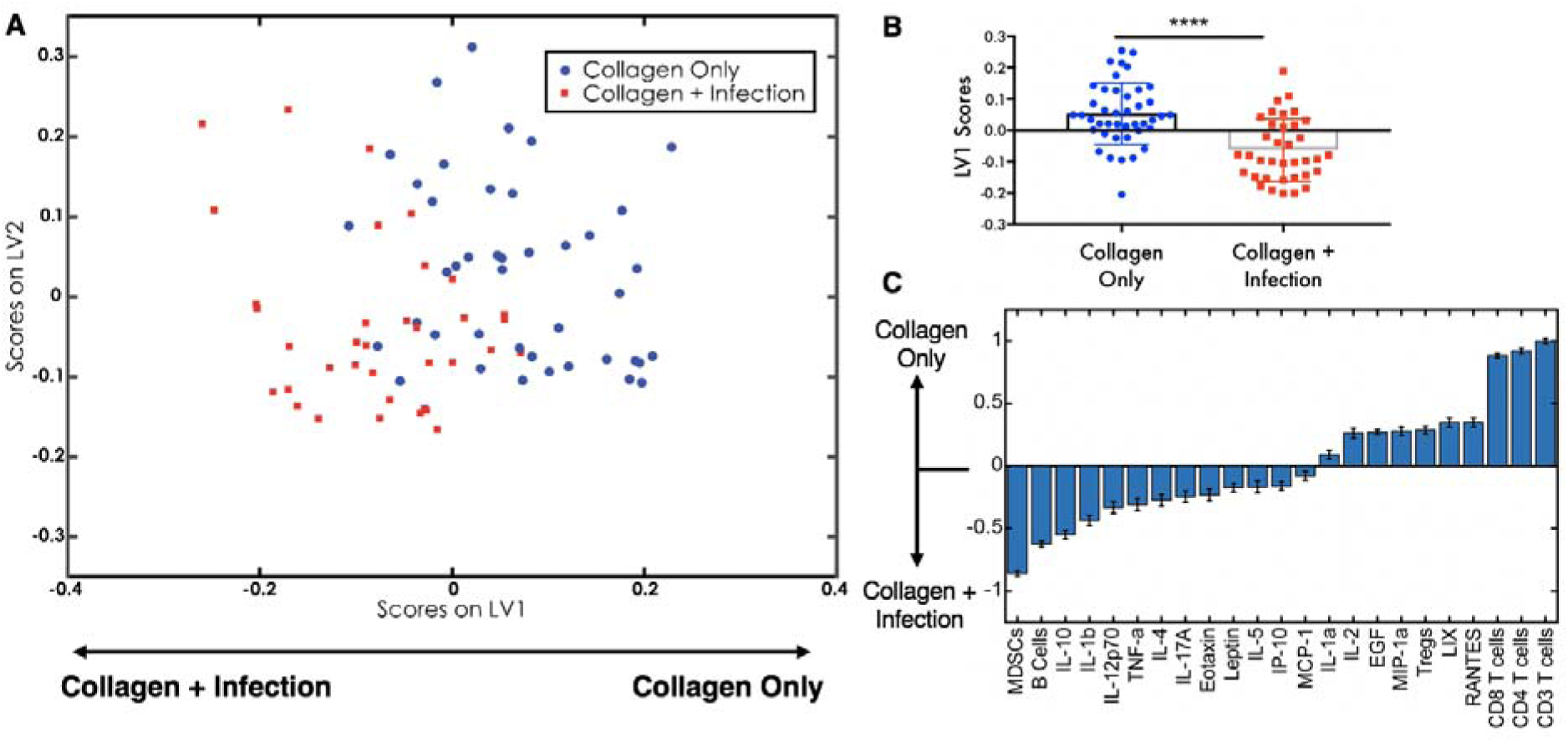
A) PLSDA plot shows all cytokine levels and cell population values post-surgery for the collagen only group (blue squares) and the collagen + infection group (red circles) plotted on latent variable 1 (LV1) and latent variable 2 (LV2) axes. B) Plotting only LV1 scores shows a significant difference (p<0.001 (****) according to Student’s t-test) between the collagen only and collagen + infection group based on LV1. C) The LV1 loading plot shows the top factors that most contributed to positive LV1 scores (right) and the top factors that most contributed to negative LV1 scores (left).

## DISCUSSION

Despite advancements in surgical procedures and antimicrobial regimens, orthopedic biomaterial-associated infections remain a challenging clinical problem with high failure rates and little to no improvement in infection-related patient outcomes over the past several decades (36). In particular, for fracture-associated infections, there is currently no consensus on diagnostic criteria and therefore very few protocols for diagnosis and treatment (37). *S. aureus* causes the majority of bone infection cases and is known for its immune evasion mechanisms and manipulation of humoral and adaptive immunity. Because the immune system is a key player for the clearance of bacterial infections and tissue regeneration, there has been an increased focus on targeted immune modulation as a potential tool for the treatment of chronic biomaterial-associated orthopedic infections (12). Despite the inextricable link between the local and systemic immune environments, there has been little focus placed on understanding the role of the systemic immune response in particular. Recent work has shown that systemic immune homeostasis is altered by local biomaterial scaffolds for tissue regeneration (38), and more interestingly, that systemic immunity is required for successful anti-tumor immune therapy, suggesting that local immune modulation alone is not sufficient for successful intervention (39). Therefore, characterization of the systemic immune response to severe trauma with biomaterial-associated infection could allow for better identification of immunotherapeutic targets that could improve local therapeutic interventions and overall outcomes for these patients. In this manuscript we utilized a rat trauma model with a biomaterial-associated infection to analyze the long-term immune response to a local, indolent orthopedic infection with the hypothesis that systemic immune dysregulation and immunosuppression would develop. Long-term systemic immune dysregulation was observed over 8 weeks (56 days) and as early as Day 1 post-injury in our rat trauma infection model with elevated levels of immunosuppressive MDSCs and decreased levels of effector T cells along with a dysregulated and uncoordinated cytokine response compared to non-infected trauma rats. This systemic immune dysregulation may further enhance immune evasion tactics of *S. aureus* and hinder successful bone regeneration in cases of orthopedic biomaterial-associated infections, although further work is needed to better understand the role of the systemic response on bacterial clearance and healing.

In the presence of orthopedic infection in our rat model of infected trauma, there was depressed systemic immunity compared to trauma without infection, most notably with increases in immunosuppressive MDSCs and decreases in T cells, including the helper T cell and cytotoxic T cell subsets. MDSCs are a heterogeneous and immature myeloid-derived cell population characterized by their immunosuppressive function, and they expand during conditions of acute and chronic inflammation, including trauma, sepsis, infection, and cancer (18,19,40,41). MDSCs utilize various mechanistic pathways to suppress the activity of immune effector cells, including the production of immunosuppressive cytokines such as IL-10 and TGF-β, release of reactive oxygen and reactive nitrogen species (ROS/RNS), and stimulation of immunosuppressive Tregs (42,43). Additionally, MDSCs can suppress T cell function through arginase-1 activity that depletes L-arginine, an essential amino acid required for T-cell receptor (TCR) signaling (44,45). Prevention of TCR and T cell activation inhibits one of the major response mechanisms to pathogen-associated molecular patterns (PAMPs) and danger-associated molecular patterns (DAMPs), allowing bacteria to evade recognition and clearance by the immune system. Additionally, in the context of infection and cancer, MDSCs are known for suppressing cytotoxic T cell and Natural Killer (NK) cell function, preventing immune responses to bacteria and tumors (41,46). Interestingly, in our study we observed that B cells were elevated in the blood in either the infected group or the non-infected group compared to each other depending on the timepoint. Initially, B cells in the infected group increased to levels higher than the non-infected group, peaking around Day 7. Following Day 7, B cell levels in the infected group notably decline, whereas B cell levels in the collagen only, non-infected group remain relatively constant. *S. aureus* is known to modulate the B cell response by triggering B cell expansion through Staphylococcal Protein A (SpA) and inducing immune tolerogenic IL-10 producing B cells. This is supported by our data showing a significant increase in IL-10 in the infected group compared to the collagen only group (47,48). Following this expansion, *S. aureus* immune evasion mechanisms subsequently lead to an ablation of B cells, in particular the long-term plasma cells, thus increasing the risk for chronic and recurring infection (49), which is supported by our data showing a continual decline in B cells in the infection group after a peak at Day 7. While there were overall differences in systemic cellular immunity between the infected and non-infected groups, there were few significant differences at individual timepoints. The lack of individual differences in cell populations may in part be due to the complexity of the interactions between the different mediators. Additionally, as a limitation of this study, we did not conduct functional analyses of the various immune cell populations or investigate further heterogeneity within subsets of cell types, which may highlight further and more significant differences in these populations.

Tissue cellular analysis showed similar results as the systemic analysis with upregulation of immunosuppressive MDSCs in the local soft tissue surrounding the defect, as well as in the bone marrow. Macrophages were elevated in the collagen only, non-infected group in the bone marrow and the spleen, but most dramatically in the local soft tissue. *S. aureus* biofilm formation can alter the local environment in order to impair immune effector cell function and enhance the expansion of immunosuppressive MDSCs (12). In the presence of infection, the immunosuppressive immune environment can prevent infiltration of macrophages, whereas, the lack of infection in the trauma only group can permit extensive macrophage infiltration into the defect site. Due to an emphasis on the systemic immune response, one limitation of this study was a lack of characterization of macrophage phenotype infiltrating into the defect region. Future work to better understand the interactions between both the local and systemic environments could be essential for identifying appropriate immunomodulatory targets to enhance bacterial clearance and improve tissue regeneration.

Overall analysis of circulating cytokine levels reveals a regulated and coordinated immune response with resolution of inflammation sometime between Day 14 and Day 28 in the collagen only, non-infected group. In contrast, the infection group lacked a similar coordinated response at any timepoint or with any set of cytokines compared to the collagen only group. Looking at significant cytokine differences between groups prior to Day 7, there was early upregulation of cytokines involved in cell recruitment, proliferation, and survival in the non-infected group compared to the infected group, including EGF, MCP-1 (also known as CCL2), and LIX (also known as CXCL5) at Day 1, and RANTES (also known as CCL5) at Day 3. Under inflammatory conditions, EGF is a potent inducer of angiogenesis and bone growth by significantly upregulating secretion of vascular endothelial growth factor (VEGF) and bone morphogenetic protein-2 (BMP-2), an osteoinductive growth factor (50). MCP-1 has also been shown to be a key cytokine involved in both inflammation and bone remodeling, in particular with processes that include neutrophil migration, macrophage infiltration, and angiogenesis (51). Similarly, LIX and RANTES are known for their chemotactic and pro-inflammatory properties that have both been associated with the necessary inflammatory phase during the fracture repair process (52,53). In the infection animals on the other hand, immunosuppressive IL-10 was upregulated as early as Day 3 compared to the non-infected animals, which may have subsequently inhibited a coordinated and regulated inflammatory response. This increase in IL-10 is supported by a sharp increase as early in MDSCs as early as Day 1 in the infection group, which are known to release IL-10. At Days 7 and 14, multiple inflammatory cytokines, including were upregulated in the collagen only group compared to the infection group, which all resolved back to baseline levels by Day 28. It has generally become accepted that a regulated inflammatory response is required and even beneficial for repair and regeneration of severe injuries in order to recruit appropriate cells and clear necrotic tissue (54). However, a similar inflammatory response is not seen in the infection group. At Day 56, several cytokines including IL-4, IL12p70, IL-17A, and TNFa are upregulated in the infection group compared to the collagen only group, highlighting that the dysregulated cytokine response continues long-term in the infection group.

In addition to looking individually at cell populations and cytokine levels, we also conducted multivariate analyses that can account for interaction effects between these different factors to allow us to better understand the role of these systemic factors in response to orthopedic biomaterial-associated infected trauma. The PLSDA loading plot showed mainly cell types, not cytokines, as the top factors associated with each group. The collagen only, non-infected group was most positively associated with T cells, including the helper T cell and cytotoxic T cell subsets. The presence of T cells demonstrates the importance of functioning immune effector cells for a coordinated and regulated immune response to trauma. In particular, an absence of T cells has been shown to result in disrupted bone mineralization and decreased bone regeneration following fracture. This effect has been associated with the necessity of T cells for normal collagen deposition during the early stages of callus formation (55,56). In addition, systemic dysregulation of helper T cells has been associated with increased risk for multiple organ failure and death in trauma and burn patients (57), again highlighting the importance of T cells and their subsets for appropriate healing. In contrast, the top factors most associated with the infection group included MDSCs, B cells, and IL-10. Immunosuppressive MDSCs, which are known to release the cytokine IL-10 and expand following bacterial infection, can be a sign of a dysregulated immune response (41). In addition, *S. aureus* is also known to modulate the B cell response by enhancing the population of immune tolerogenic IL-10 producing B cells (47,48).

Although differences in the immune response were observed between the two groups, no significant differences in bone regeneration volumes were observed, despite the sub-critical size of the bone defect. In this study, bone defects in the non-infected group were treated with collagen sponge, which is clinically approved for use in patients in conjunction with BMP-2, an osteoinductive growth factor. Although our treatment strategy did not use BMP-2, collagen is one of the most prevalent molecules in the extracellular matrix (ECM) and has had success in tissue engineering and wound healing applications as a scaffold to facilitate cell growth and differentiation. Collagen gels used for growth factor and cell delivery have been shown to promote intrinsic vascularization and provide an infrastructure for osteogenesis and chondrogenesis by supporting cell invasion (58,59). While there are many collagen-based scaffolds and hydrogels, only bovine collagen I scaffold (collagen sponge) is approved for clinical use in bone regeneration applications in combination with BMP-2. However, the effects of this particular scaffold alone on bone healing remain unknown since typically BMP-2 is co-delivered on the scaffold. In a recent study, *in vivo* testing of collagen sponge unexpectedly showed a negative impact on bone healing, with significantly decreased bone formation in a mouse osteotomy model compared to an empty defect (60). These results are consistent with the lack of healing observed in our study for the non-infected, collagen only treatment group; however, further mechanistic studies are needed to understand the potential inhibitory effects of empty collagen sponge in sub critical-size bone defects.

*S. aureus-*soaked collagen sponges were implanted within the defect site in order to create a local, indolent infection associated with an orthopedic biomaterial. Animals did not develop a fever or lose weight compared to non-infected animals, indicating that the infection remained local and did not spread systemically. To confirm bacterial presence in the infection animals, longitudinal bioluminescent imaging and endpoint wound swab culture were conducted. Despite a lack of bioluminescent signaling past Day 7, culture of the wound swab at the 8-week endpoint confirmed the presence of bacteria. More recently, it has been proposed that *S. aureus* biofilm formation results in physiologically dormant cells that are protected by the biofilm matrix, contributing to antibiotic resistance (61,62). Additionally, these dormant “persister” cells are thought to grow within the biofilm at higher rates than metabolically active planktonic cultures (63). Therefore, the lack of bioluminescent signal past Day 7 could be indicative of this biofilm formation and a transition of these cells to a less metabolically active state.

This is one of the first studies investigating the systemic immune response to orthopedic biomaterial-associated infection following trauma. The presence of a local, indolent *S. aureus* bacterial infection resulted in an uncoordinated and dysregulated systemic immune response, with systemic increases in immunosuppressive MDSCs and decreases in immune effector cells, including T cells. This systemic immune dysregulation and immunosuppression further exacerbated local immune evasion techniques by *S. aureus*, which could contribute to the clinical challenges associated with infected trauma, in particular, chronic and recurring infections. An improved understanding of the systemic immune response and its relationship with the local environment, including bacterial clearance and bone healing, could provide promising systemic immunomodulatory targets to improve the ability to fight implant-associated infections and improve patient outcomes.

The data from this study indicate that a local, indolent orthopedic biomaterial-associated infection has widespread systemic effects, in particular, on the immune response. Characterization of the long-term systemic immune response to the biomaterial-associated infection group revealed evidence of systemic immune dysregulation and immunosuppression, with increased immunosuppressive MDSCs and decreased immune effector T cells. Further, the systemic cytokine response in the presence of infection appeared dysregulated and lacked coordination compared to the systemic cytokine response in the absence of infection. The presence of infection also resulted in decreased macrophage infiltration in the surrounding soft tissue at 8 weeks post-injury and infection, highlighting the complicated immune environment that results from orthopedic-associated infections. Further work will be essential to better understand the development of systemic immune dysregulation and its role on bacterial clearance and bone defect healing.

## ACKNOWLEDGEMENTS

This research was supported by an ACTSI/REM Seed Grant from Emory University and Georgia Tech. CEV acknowledges support by the National Science Foundation Graduate Research Fellowship under Grant No. DGE-1650044. Any opinion, findings, and conclusions or recommendations expressed in this material are those of the authors(s) and do not necessarily reflect the views of the National Science Foundation. Additionally, the authors wish to acknowledge the core facilities at the Parker H. Petit Institute for Bioengineering and Bioscience at the Georgia Institute of Technology for the use of their shared equipment, services, and expertise. And to thank Laura Weinstock for her assistance with the Luminex analysis.

