## Supplemental Data for "Development of Systemic Immune Dysregulation in a Rat Trauma Model with Biomaterial-Associated Infection"

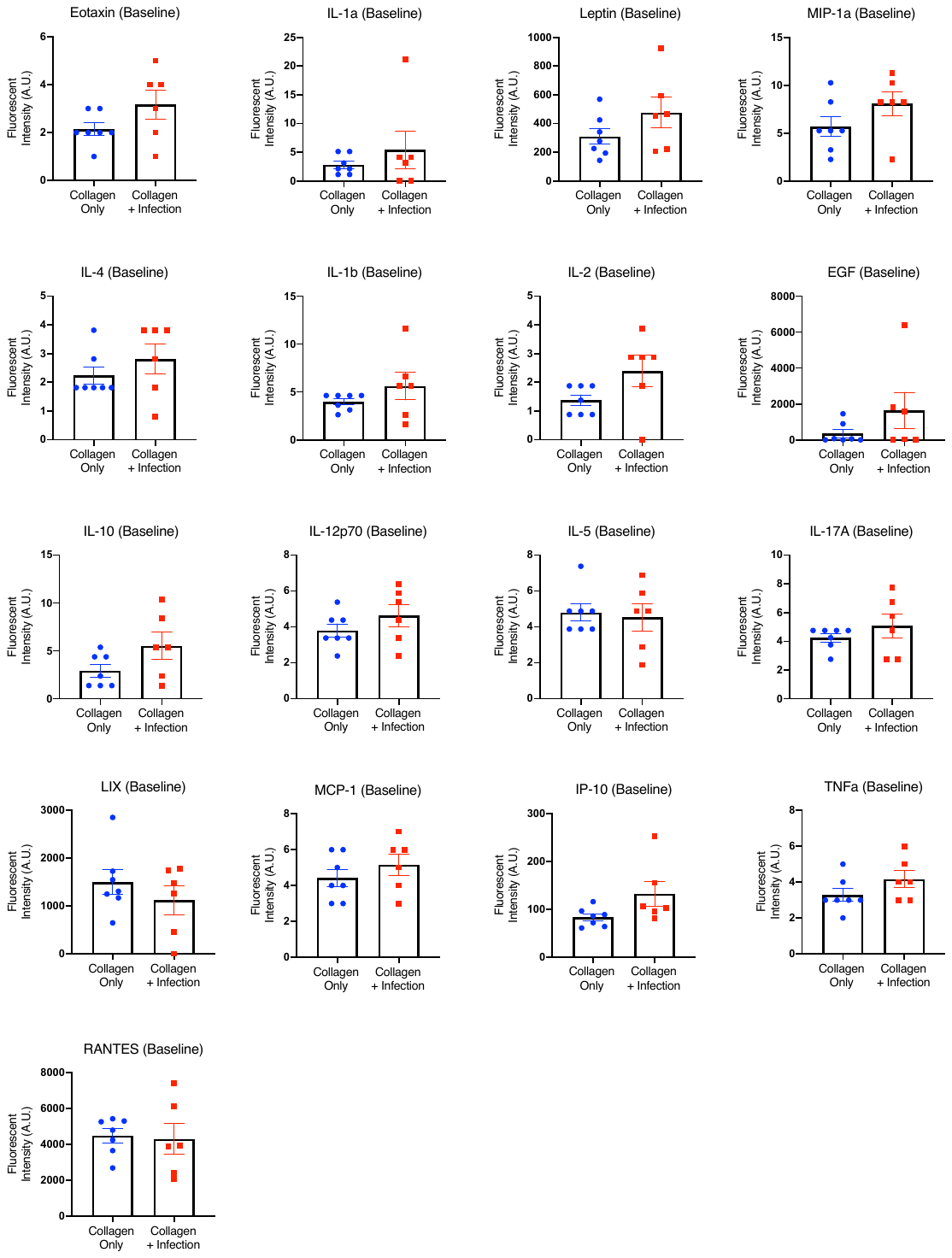

**Supplementary Figure 1**

**A**

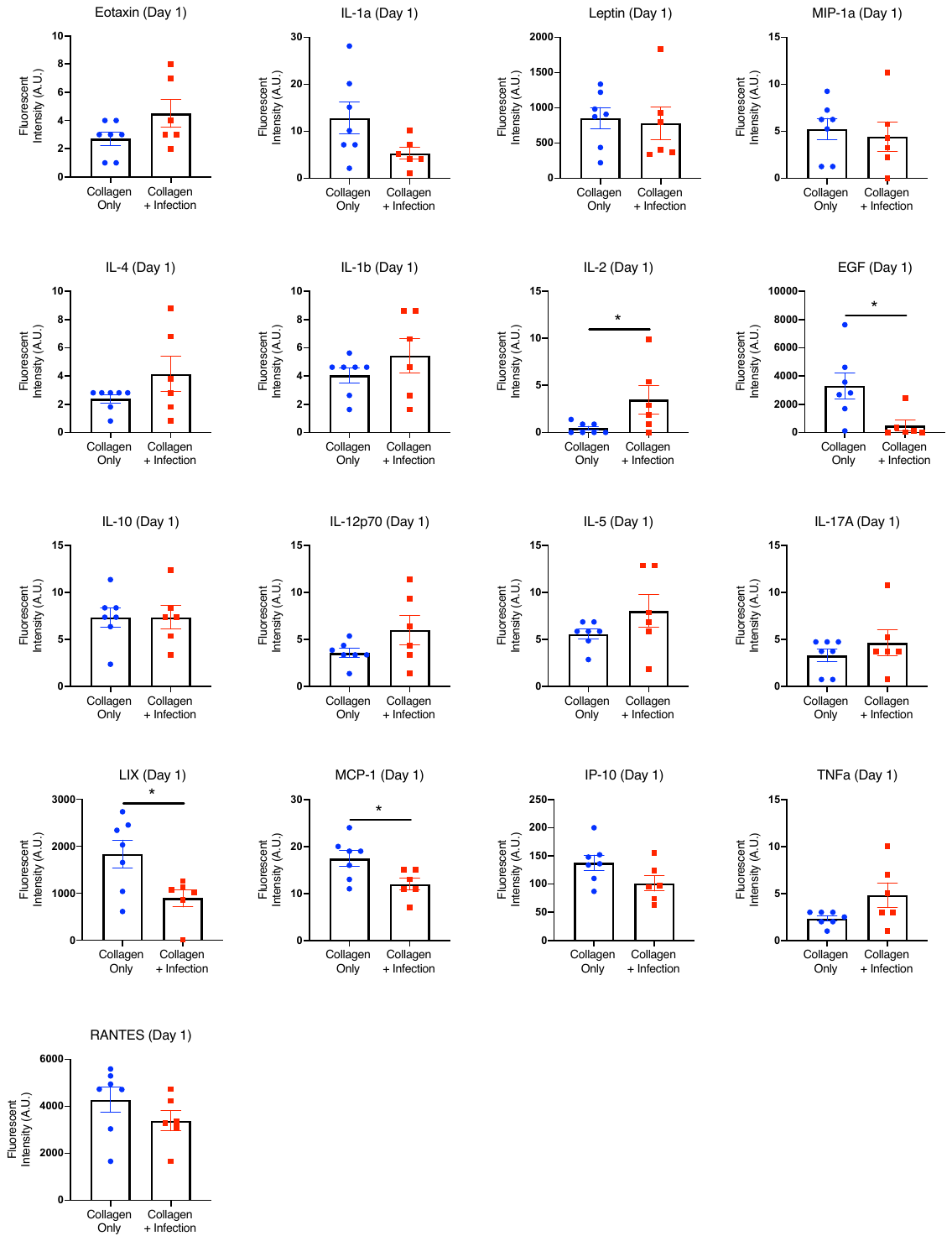

**B**

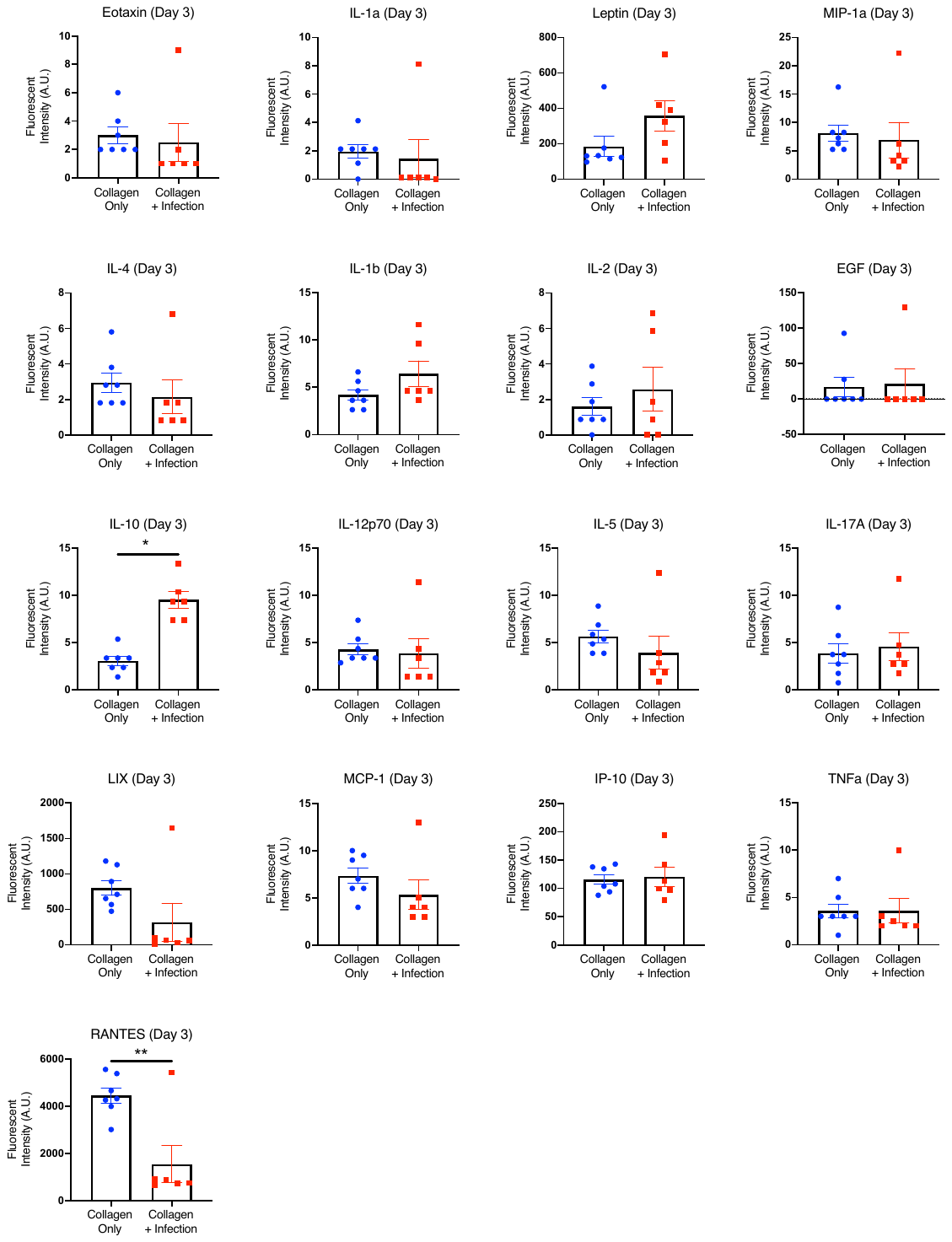

**C**

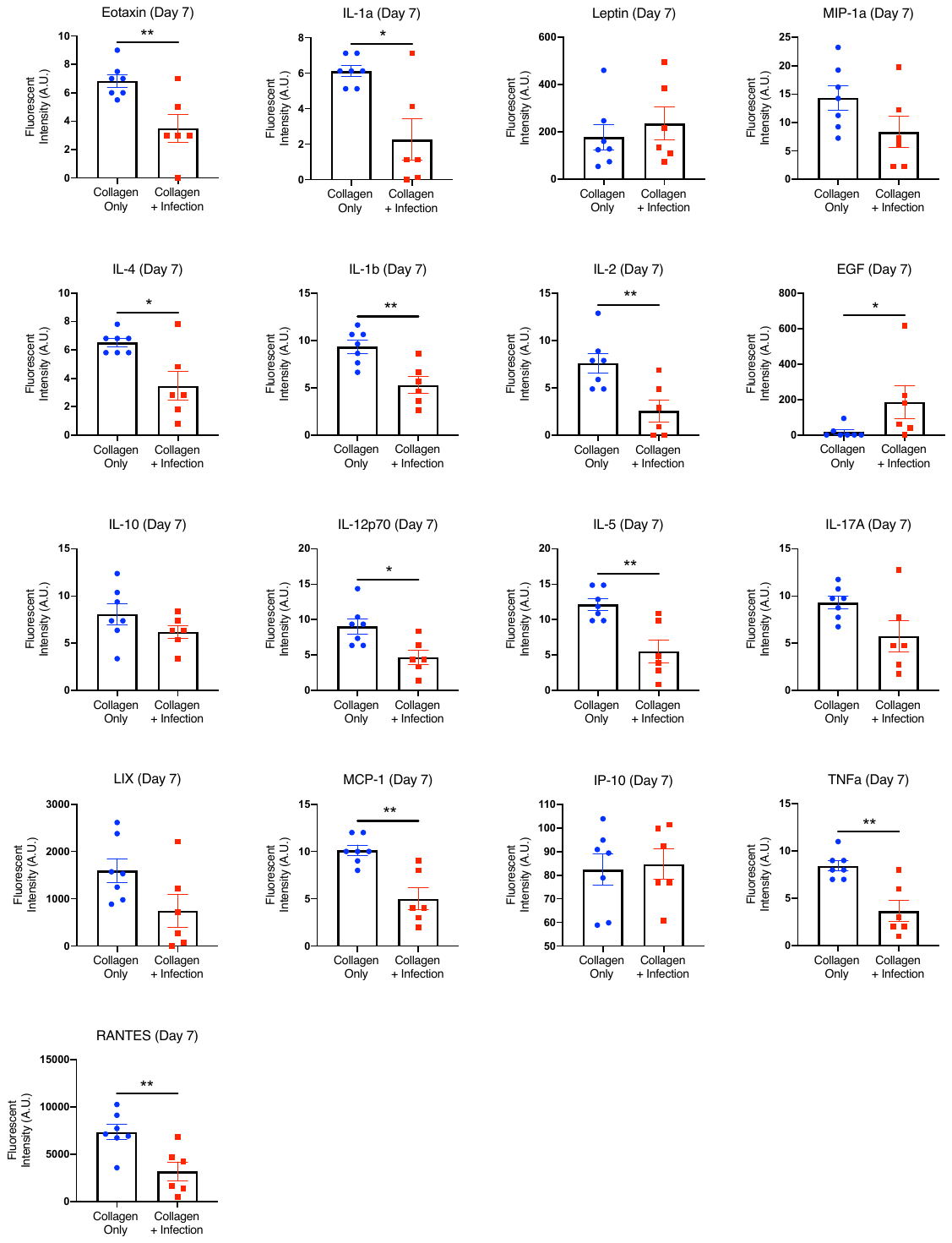

**D**

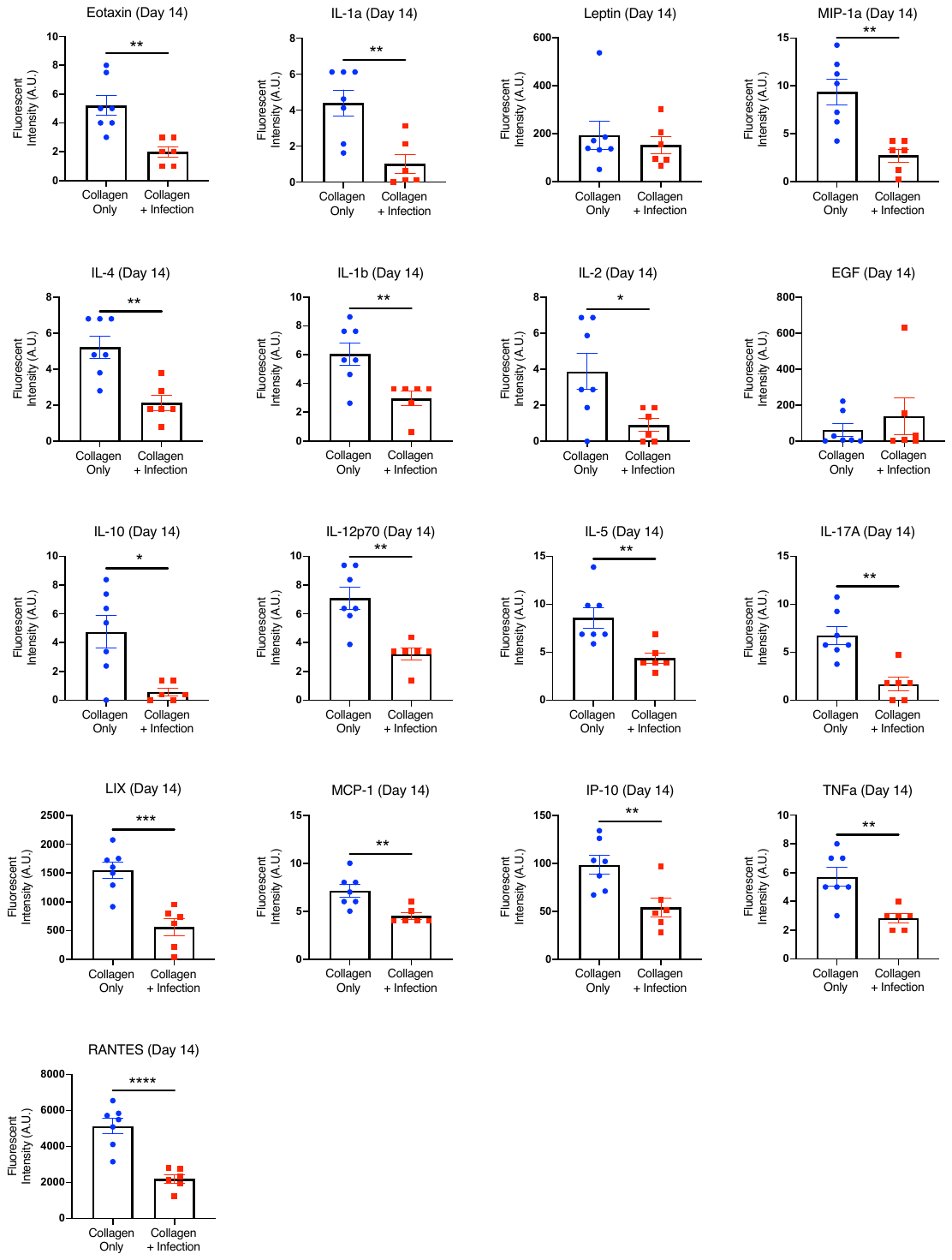

**E**

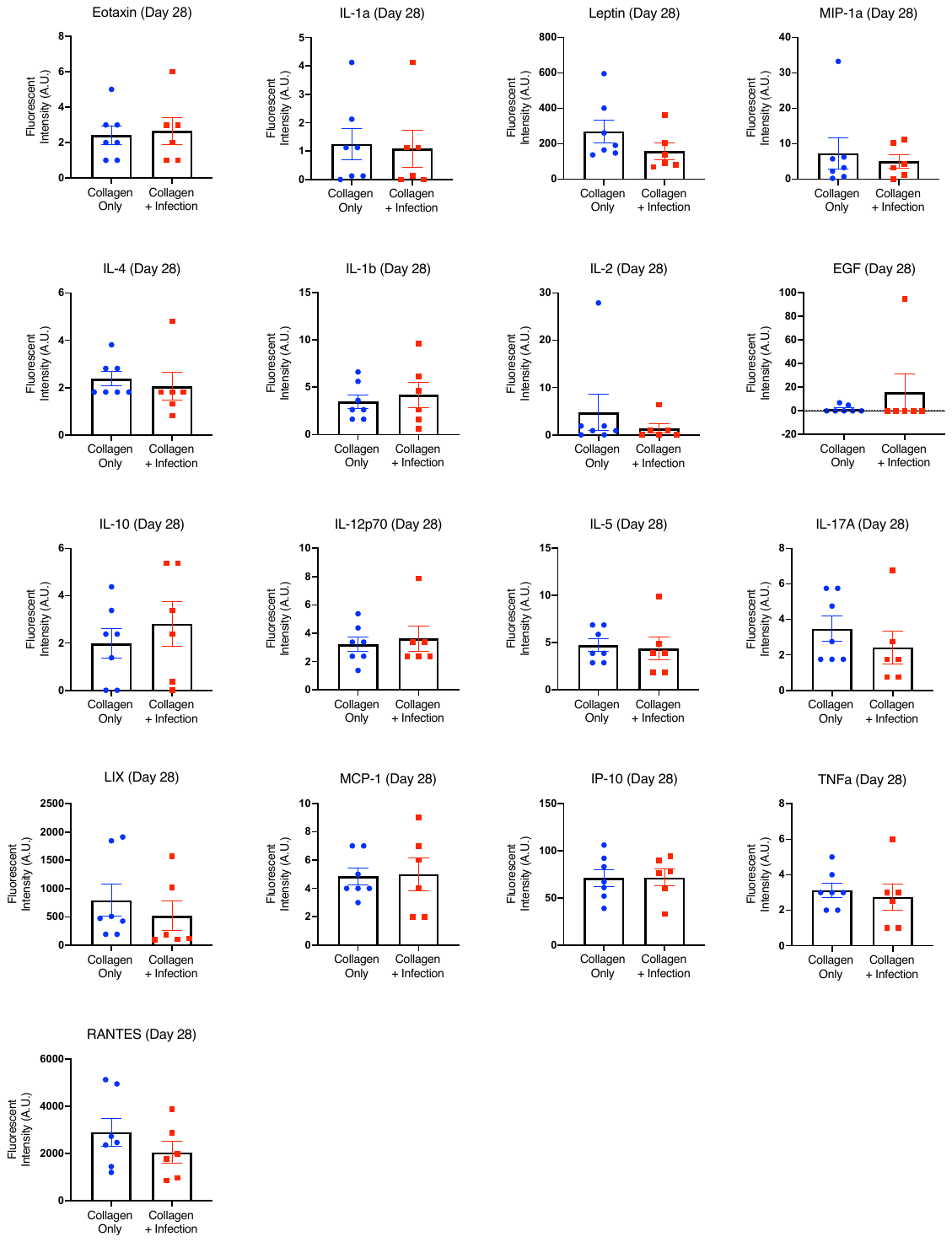

**F**

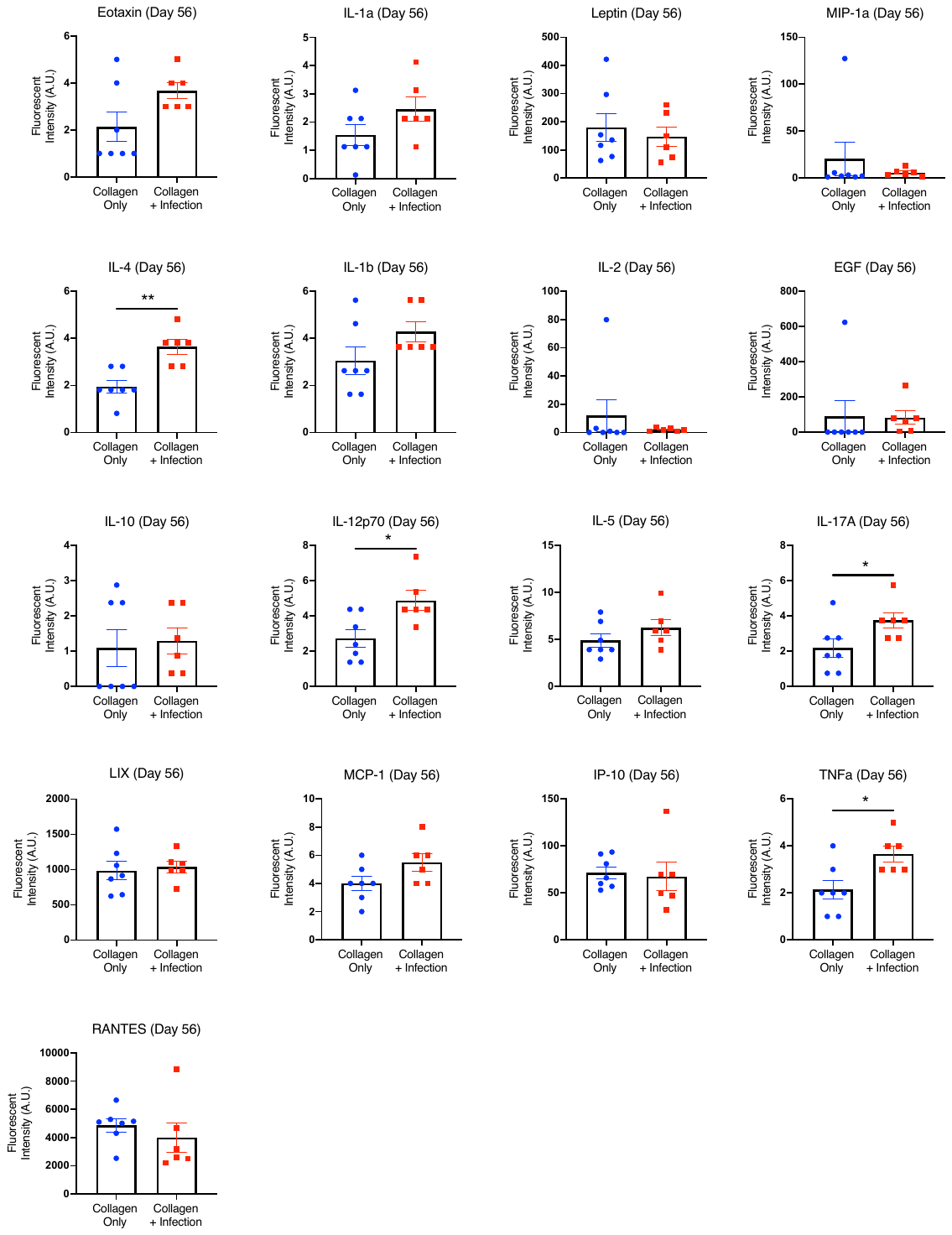

**G**

**Supplementary Figure 1.** Cytokine levels in the collagen only and infection groups at the baseline (A) and at Day 1 (B), Day 3 (C), Day 7 (D), Day 14 (E), Day 28 (F), and Day 56 (G). Significance was determined using Student’s t-test or with Mann-Whitney test for non-parametric data with p<0.05 (*), p<0.01 (**), p<0.005 (***), and p<0.001 (***).
